# 6-Thioguanine blocks SARS-CoV-2 replication by inhibition of PLpro protease activities

**DOI:** 10.1101/2020.07.01.183020

**Authors:** Caleb D. Swaim, Yi-Chieh Perng, Xu Zhao, Larissa A. Canadeo, Houda H. Harastani, Tamarand L. Darling, Adrianus C. M. Boon, Deborah J. Lenschow, Jon M. Huibregtse

**Affiliations:** Department of Molecular Biosciences, University of Texas at Austin, Austin, TX; Department of Internal Medicine, Washington University School of Medicine, St. Louis, MO; Department of Pathology and Immunology, Washington University School of Medicine, St. Louis, MO; Department of Molecular Microbiology, Washington University School of Medicine, St. Louis, MO

## Abstract

A recently emerged betacoronavirus, SARS-CoV-2, has led to a global health crisis that calls for the identification of effective therapeutics for COVID-19 disease. Coronavirus papain-like protease (PLpro) is an attractive drug target as it is essential for viral polyprotein cleavage and for deconjugation of ISG15, an antiviral ubiquitin-like protein. We show here that 6-Thioguanine (6-TG) inhibits SARS-CoV-2 PLpro-catalyzed viral polyprotein cleavage and ISG15 deconjugation in cells and inhibits replication of SARS-CoV-2 in Vero-E6 cells and Calu3 cells at submicromolar levels. As a well-characterized FDA-approved orally delivered drug, 6-TG represents a promising therapeutic for COVID-19 and other emerging coronaviruses.

**One Sentence Summary:** A repurposed drug that targets an essential enzymatic activity of SARS-CoV-2 represents a promising COVID-19 therapeutic.

## Main Text

Coronavirus Disease-2019 (COVID-19) is caused by SARS-CoV-2, a betacoronavirus highly similar to SARS (now SARS-CoV-1) (*1, 2*). An urgent need exists for treatment strategies, and repurposed FDA-approved drugs with existing supply chains and well-characterized pharmacologic properties represent a rapid and efficient approach toward COVID-19 therapeutics (*3*).

Enzymes encoded by SARS-CoV-2 are attractive drug targets, including PLpro, an essential cysteine protease activity within the nsp3 protein (*4*). PLpro cleaves the pp1a polyprotein at three sites, generating mature nsp1, 2 and 3 proteins. The PLpro recognition site (LXGG) is also found at the C-terminus of ubiquitin and ISG15, an antiviral ubiquitin-like modifier, although SARS-CoV-1 and -2 PLpro preferentially catalyze de-ISGylation over de-ubiquitylation (*5, 6*). Therapeutic inhibition of PLpro would therefore be predicted to have two antiviral effects: restoration of the antiviral effect of ISGylation and inhibition of viral replication by blocking polyprotein cleavage. Further, de-ISGylation by PLpro, through generation of free (unconjugated) ISG15, enhances the secretion and extracellular signaling function of ISG15, which in turn promotes pro-inflammatory cytokine production from cells of the immune system (*7*). Therefore, a potential third effect of inhibiting PLpro would be a decrease in pro-inflammatory “cytokine storms” associated with COVID-19 (*8*).

6-Thioguanine (6-TG) is an FDA-approved drug that has been used in the clinic since the 1950s, originally for the treatment of childhood leukemias and subsequently for treatment of inflammatory bowel and Crohn’s disease (*9*). 6-TG was previously reported to inhibit the SARS-CoV-1 and MERS PLpro catalytic domain *in vitro* (*10, 11*), with an IC50 of 5 µM and 24.4 µM, respectively, however there was no further follow up of its ability to inhibit de-ISGylation or viral polyprotein cleavage in cells or its ability to inhibit viral replication. Fig. 1A shows that co-expression of PLpro (residues 746-1061 of nsp3) with ISG15 and its conjugation enzymes (Uba7, Ube2L6, and Herc5) in HEK293T cells resulted in nearly complete loss of ISG15 conjugates, while expression of the active-site C-to-A variant of PLpro (C856 of nsp3; PLpro^CA^) was inactive. Addition of 6-TG resulted in a dose-dependent inhibition of PLpro de-ISGylation activity, with half-maximal inhibition at approximately 0.1 µM (Fig. 1A; quantitation shown in Supplementary Fig. 1A).

**Figure 1.**
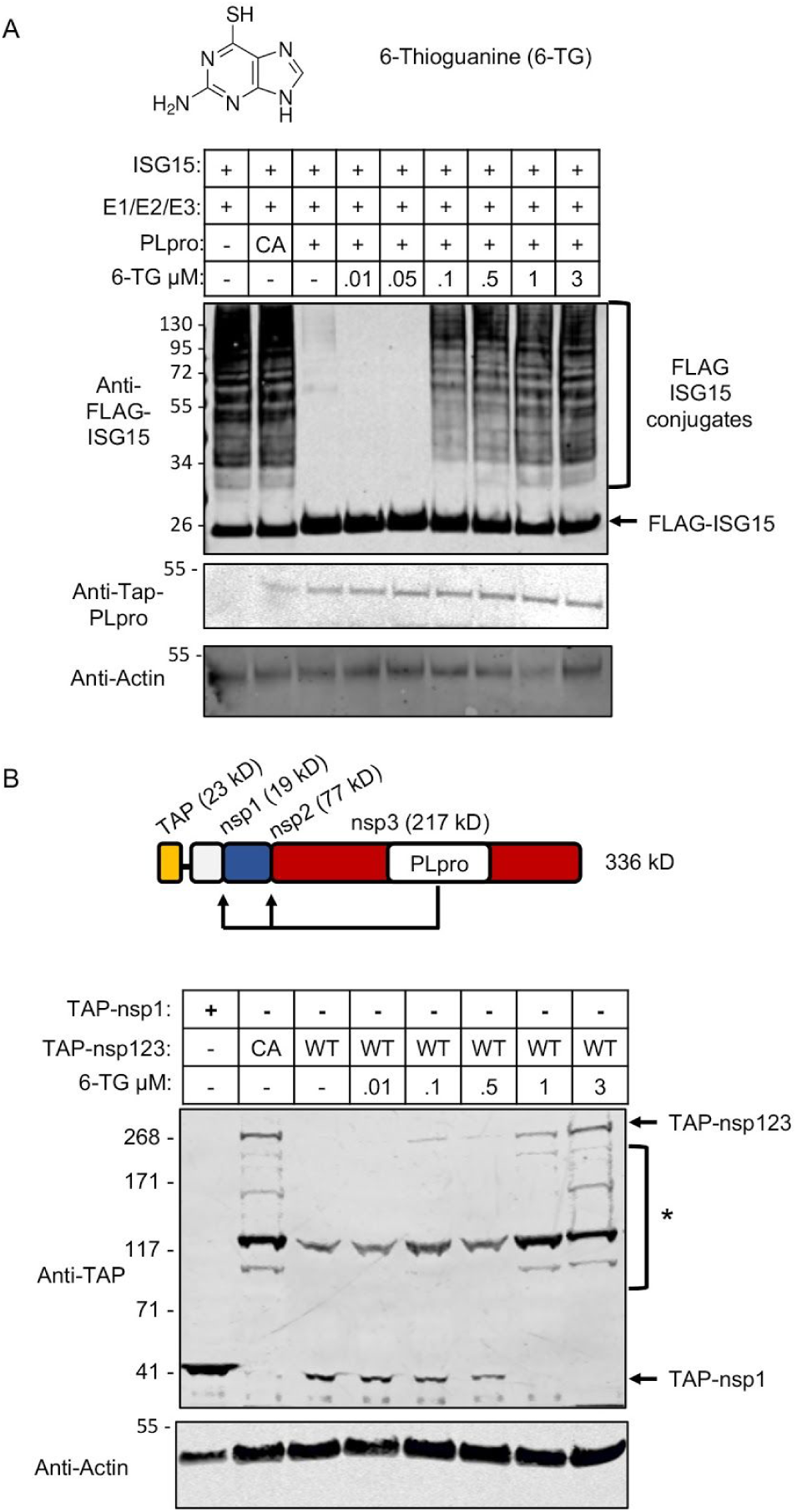
6-TG inhibits PLpro mediated de-ISGylation and viral polyprotein cleavage. **A.** The structure of 6-Thioguanine is shown. HEK293T cells were transfected with plasmids expressing FLAG-ISG15 and the ISG15 E1/E2/E3 enzymes and 0.2 μg of PLpro plasmid (WT or CA) where indicated. 6-TG was added at the time of transfection at the indicated final concentrations (μM). Total cell lysates were prepared 48 hours post-transfection and analyzed by immunoblotting to detect FLAG-ISG15 and FLAG-ISG15 conjugates. **B.** (Top) Schematic representation of the TAP-nsp123 fusion protein, with molecular masses of subdomains indicated. Arrows indicate sites of PLpro-catalyzed cleavage. (Bottom) HEK293T cells were transfected with plasmids expressing either TAP-nsp1, TAP-nsp123^WT^ or TAP-nsp123^CA^. 6-TG was added at the time of transfection at the indicated final concentrations (μM) and total cell lysates were prepared 48 hours post-transfection and analyzed by immunoblotting with anti-TAP antibody. Bands corresponding to TAP-nsp1 and the full-length fusion proteins are indicated. Bracketed bands (*) represent breakdown products of the full-length WT or CA fusion proteins.

The PLpro domain, located within nsp3, generates mature nsp-1, 2 and 3 proteins from the pp1a polyprotein through self-catalyzed cleavage reactions (Fig. 1B, schematic). We determined whether 6-TG inhibited polyprotein processing, as this is an essential function of Plpro and the natural context in which PLpro is expressed. An N-terminally TAP-tagged fusion of nsp1, 2 and 3 (TAP-nsp123^WT^) was expressed in HEK293T cells (Fig. 1B, schematic), as well as the active-site C-to-A variant (TAP-nsp123^CA^). The fusion protein was detected with anti-TAP antibody, and TAP-nsp1 was expressed separately as a size marker for the fully processed fusion protein. As shown in Fig. 1B, expression of TAP-nsp123^WT^ resulted primarily in a product the size of TAP-nsp1 (42 kD), as expected if the fusion protein was fully processed, while expression of TAP-nsp123^CA^ resulted in the expected full-length protein (∼336 kD) along with several breakdown products of the full-length protein. Increasing concentrations of 6-TG inhibited PLpro-mediated processing of the TAP-nsp123^WT^ polyprotein, with loss of the TAP-nsp1 product and accumulation of the full-length fusion protein, with the same pattern of breakdown products seen with the TAP-nsp123^CA^ protein. Half-maximal inhibition of polyprotein cleavage occurred between 0.1 and 0.5 µM (quantitation shown in Supplementary Fig. 1B). This result indicates that 6-TG inhibits an essential function of PLpro activity and that it inhibits PLpro when it is expressed in the context of the full-length nsp3 protein.

In addition to its function as a ubiquitin-like modifier, free (unconjugated) ISG15 exits cells and signals to LFA-1-expressing cells of the immune system (*e.g*., NK cells, T cells) to release specific cytokines, including IFN-γ and IL-10 (*7, 12*), while ISG15 conjugates are retained in cells. Therefore, a secondary effect of PLpro-mediated de-ISGylation is the enhanced secretion of ISG15 (*7*). We determined whether 6-TG would block this effect. Fig. 2A shows that expression of PLpro in HEK293T cells led to decreased intracellular ISG15 conjugates (upper blot, total cell lysates) and increased free ISG15 in the extracellular space (lower blot, IP-western of ISG15 from cell culture supernatants), and that 6-TG not only restored intracellular conjugates but also blocked the production of free extracellular ISG15.

**Figure 2.**
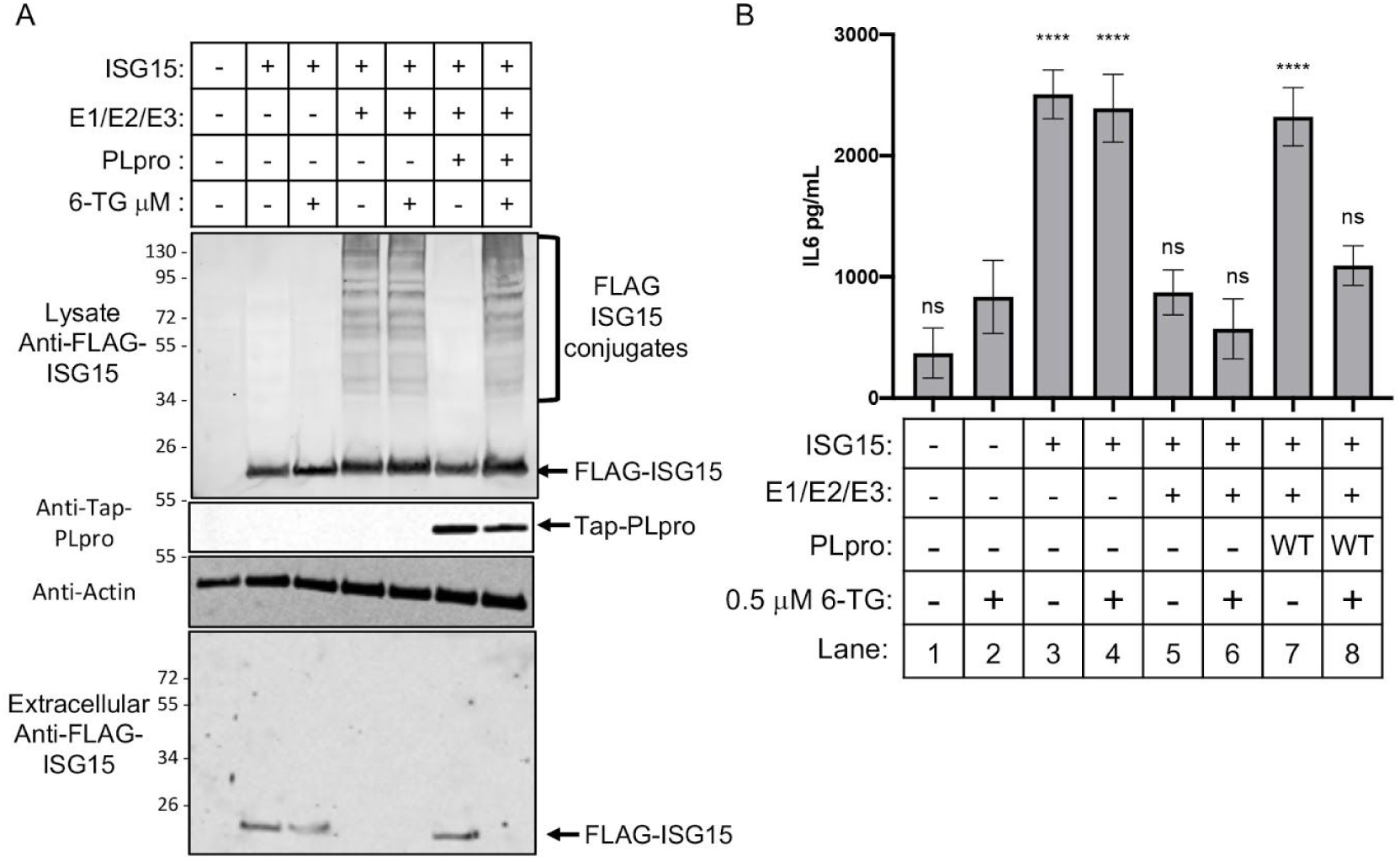
6-TG inhibits PLpro-enhanced extracellular ISG15 signaling and production of IL-6 from PBMCs. **A.** HEK293T cells were transfected and treated with 6-TG, as indicated. Total cell lysates were prepared and cell culture supernatants were collected 48 hours post-transfection. The upper panel shows a FLAG-ISG15 immunoblot analysis of cell lysates and the lower panel shows a FLAG-ISG15 immunoblot of an anti-FLAG immunoprecipitation of the cell culture supernatants. **B.** HEK293T cells were plated in the lower chamber of a transwell plate and transfected with plasmids expressing FLAG-ISG15, the ISG15 E1/E2/E3 enzymes, and PLpro, as indicated. 6-TG (0.5 µM) was added to the cells at the time of transfection. PMBCs were added to the upper chamber of the transwell plate; supernatants were collected from the upper chamber 24 hours post-transfection and assayed for IL-6 by ELISA. Significance was assessed by ordinary one-way ANOVA comparison to treatment with 6-TG alone. Asterisks indicate p-values: ****<0.0001, and non-significant changes are indicated by ns.

IFN-γ, IL-10 and IL-6 are associated with uncontrolled cytokine release syndrome and have been shown to be elevated in COVID-19 patients (*13, 14*). The complete range of cytokines affected by extracellular ISG15 signaling is unknown, but the relationship of ISG15 to IFN-γ and IL-10 (*12*) led us to test whether ISG15 also induced the secretion of IL-6 from human PBMCs. In a trans-well culture system, HEK293T cells (lower chamber) were transfected with ISG15, with or without the ISG15 conjugation enzymes (E1/E2/E3) and/or PLpro. PBMCs were placed in the upper chamber, separated from the lower chamber by a 0.4 micron membrane. In this system, ISG15 released from cells in the lower chamber signals to LFA-1-expressing cells in the upper chamber, and cytokine release in the upper chamber is measured by ELISA (*7*). Fig. 2B shows that expression of ISG15, alone, in the HEK293T cells led to secretion of IL-6 from PBMCs, and this effect was diminished when the conjugation enzymes were co-expressed with ISG15 (compare lanes 1, 3, and 5). De-ISGylation catalyzed by PLpro resulted in enhanced IL-6 secretion (compare lanes 5 and 7), and the addition of 6-TG reversed the effect of PLpro, leading to diminished IL-6 production (compare lanes 7 and 8). These results identify IL-6 as an additional cytokine that is responsive to extracellular ISG15 signaling and suggest that PLpro may be at least partially responsible for enhanced cytokine responses in COVID-19 patients. Therapeutically, 6-TG inhibition of PLpro might therefore have the additional benefit of limiting excessive cytokine responses associated with poor patient outcomes.

6-TG was next analyzed for its ability to inhibit SARS-CoV-2 replication (strain 2019 n-CoV/USA_WA1/2020; *18*) in Vero-E6 African green monkey kidney cells and in the Calu3 human lung epithelial cell line. As shown in Fig. 3, 6-TG inhibited viral replication in Vero-E6 cells with a half-maximal effective concentration (EC50) of 0.647 ± 0.374 μM. By comparison, Remdesivir inhibited SARS-CoV-2 replication in Vero-E6 cells similarly, with an EC50 of 0.77 µM (*15*). 6-TG inhibited virus replication in Calu3 cells at a lower EC50, 0.061 ± 0.049 μM. While the basis of the difference between the two cell lines is not known, cell lines can vary significantly with respect to thiopurine uptake and metabolism (*16*). 6-TG did not elicit significant cellular toxicity in either Vero-E6 or Calu3 cells (CC50 >50 μM), yielding selectivity indices (SI) of >77 in Vero-E6 cells and >819 Calu3 cells. Together, the results presented here indicate that 6-Thioguanine is a direct-acting SARS-CoV-2 antiviral, inhibiting PLpro de-ISGylation, polyprotein cleavage, and viral replication at submicromolar concentrations.

**Figure 3.**
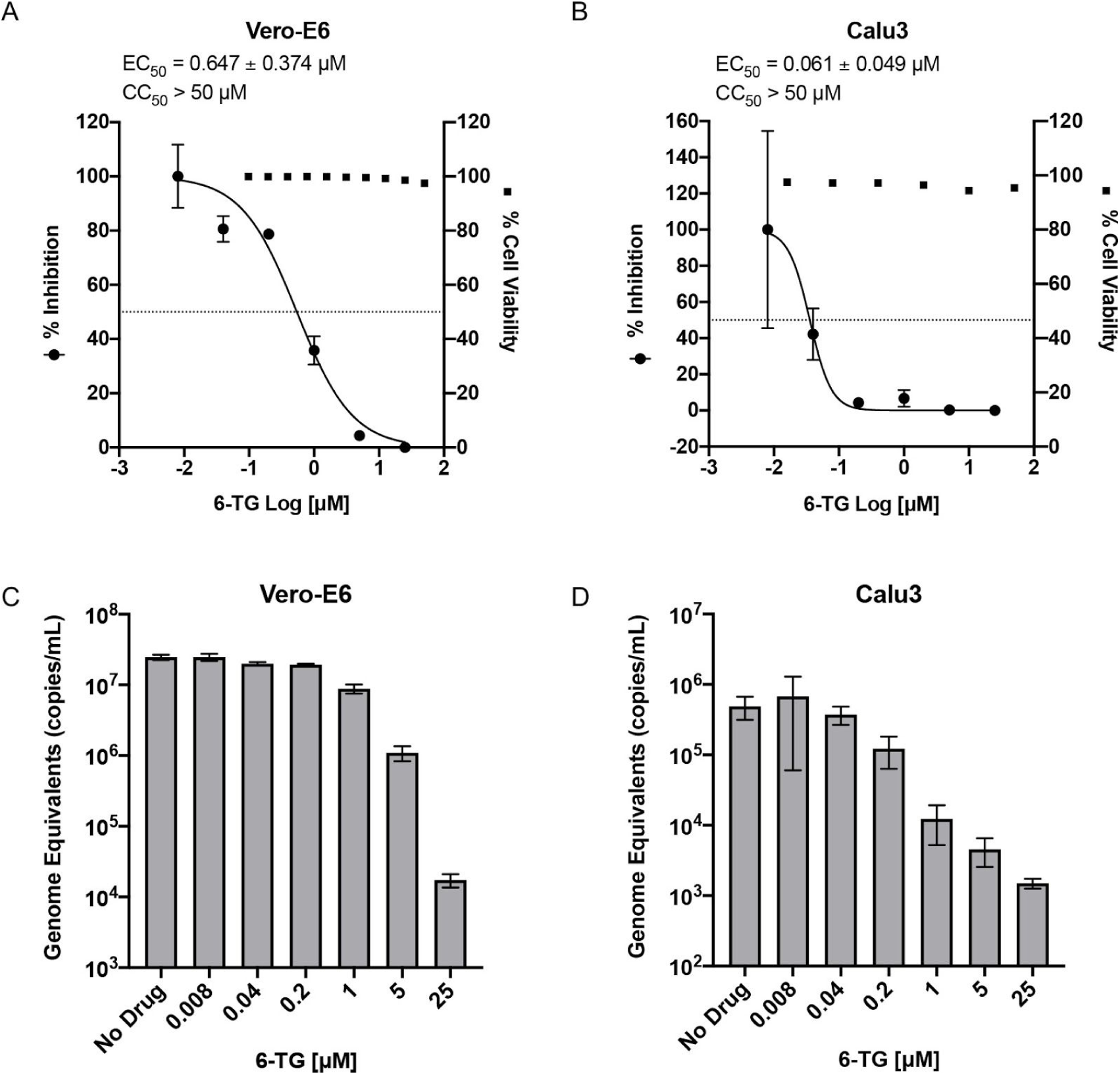
6-TG inhibits SARS-CoV-2 viral replication. Vero-E6 (**A, C**) and Calu3 (**B, D**) cells were treated with various concentrations of 6-TG and assessed for viability or were infected with SARS-CoV-2 at a multiplicity of infection (MOI) of 0.001. **A and B:** Cellular cytotoxicity of 6-TG was determined using a real-time SYTOX assay 48 hours post-treatment (closed squares) and the median cytotoxic concentration (CC50) was >50 mM in both cell lines. Inhibition of viral replication was assayed by SARS-CoV-2 N1 RT-qPCR of cell supernatants 48 hours post-infection (closed circles) and the effective median concentration for inhibition (EC50) was determined. The left and right y-axis of each graph represent mean percent inhibition of virus yield/titer and mean percent cell viability at the indicated concentration of 6-TG. **C. and D.** Viral genome equivalents in the cell supernatants were quantified by SARS-CoV-2 N1 RT-qPCR. Data are means ± SED; n = 3 – 5 biological replicates.

To our knowledge, 6-TG is the first identified FDA-approved inhibitor of PLpro demonstrated to inhibit SARS-CoV-2 replication. 6-TG is a widely available orally delivered generic drug on the World Health Organization list of essential medicines (*17*). Dosing regimens vary significantly depending on its use, from 10 mg per day for long-term treatment of inflammatory diseases, to up to 3 mg/kg/day in acute lymphocytic leukemia treatments (*18*). While toxicity can be significant at higher dosages, it is anticipated that its use as an antiviral would be over a relatively short time period and that toxicity issues would likely be minimal. We propose that the results presented here warrant the initiation of human clinical trials of 6-TG as a SARS-CoV-2 therapeutic. As PLpro is a conserved and essential enzymatic activity of the beta coronaviruses, 6-TG may prove useful in the current and future coronavirus pandemics and as a complement to other antivirals in development.

## Acknowledgments

We thank Robert Krug and Sylvie Beaudenon-Huibregtse for critical reading of the manuscript and Bill Matsui for helpful discussions.

## Author contributions

Conceptualization and methodology: J. M. H., D. J. L., C. D. S., Y.-C. P., L. A. C.; Investigation and data analysis: C. D. S., Y.-C. P., X. Z., H. H. H., T. L. D.; Supervision: J. M. H., D. J. L., A. C. M. B.; Writing, original draft: J. M. H.; Writing, editing: D. J. L., A. C. M. B., C. D. S., Y.-C. P., L. A. C.; Project administration and funding acquisition: J. M. H., D. J. L., A. C. M. B.

## Data and materials availability

All materials that are not commercially available or available from public resources are available from the corresponding author. All data is available in the main text or the supplementary materials.

## List of Supplementary Materials

Materials and Methods

Figures S1

## Supplementary Materials for

### Materials and Methods

#### Cells, viruses, and compounds

All cell lines were maintained at 37°C, 5% v/v CO_2_ in a humidified incubator. HEK293T cells were grown in DMEM (Corning) supplemented with 10% FBS (Sigma) and 1% Penicillin-Streptomycin (Corning). PBMCs from healthy individuals were isolated from Leukoreduction System (LRS) Chambers obtained from We Are Blood (Austin, TX). The LRS chamber blood was diluted 1:1 with PBS with 2% FBS and transferred to a SepMate tube (Stem Cell Technologies) containing an equal volume of Ficoll Plaque (GE Healthcare). The tube was centrifuged at 1,200 x *g* for 10 minutes at room temperature. The top layer containing the enriched mononuclear cells (MNCs) was poured off to a separate tube. The enriched MNCs were washed twice with PBS containing 2% FBS. To eliminate any red blood cells (RBCs) present in the MNC fraction, cells were lysed with 1X RBC lysis buffer (Biolegend). The MNCs were then counted and either frozen in RPMI 1640 containing 20% FBS and 5% DMSO or plated in RPMI 1640 at 1×10^6^ cells per mL.

Vero-E6 (ATCC), Calu3 (ATCC) were cultured at 37°C, 5% v/v CO_2_ in a humidified incubator in Dulbecco’s Modified Eagle medium (Corning) supplemented with 10% FBS (HyClone), 1% Penicillin-Streptomycin (Corning), 10 mM HEPES (Corning), and 1X non-essential amino acids (Sigma). Infections and culture post infection was conducted in DMEM with 2% FBS, 1% Penicillin-Streptomycin, 10mM HEPES, and 1X non-essential amino acids.

SARS-CoV-2 strain 2019 n-CoV/USA_WA1/2020 was originally obtained from the Centers for Disease Control and Prevention (CDC/BEI Resources NR52281) and was a gift of Michael S. Diamond, Washington University School of Medicine. All work with infectious SARS-CoV2 was performed in Institutional Biosafety Committee approved A-BSL3 facilities at Washington University School of Medicine using appropriate positive pressure air respirators and protective equipment.

#### Plasmids

pcDNA3.1-TAP-PLpro^WT^ and pcDNA3.1-TAP-PLpro^CA^ were described in (*7*). pcDNA3.1-TAP-nsp123 was generated from a partial orf1ab clone (S2-A2_p57 plasmids) encoding residues 1-1504; nsp1, nsp2, and part of nsp3; (gift from Hongbing Jiang and David Wang, Washington University School of Medicine) with the remainder of the nsp3 ORF (residues 1505 - 2767 of pp1a, derived from Addgene cat# 141257). The pcDNA3.1-TAP-nsp123 active site variant (C to A substitution, residue 1677 of polyprotein 1a, corresponding to residue 856 of mature nsp3) was generated from this clone by overlapping PCR. pGEX6P ISG15 was described previously (*12*) and the pGEX6P ISG15-precursor was made by adding nucleotides corresponding to the amino acid sequence TEPGGRS YPYDVPDYA (the natural ISG15 C-terminal extension and an HA tag) to the 3’ end of pGEX6P ISG15. Plasmids expressing FLAG-ISG15, and the ISG15 E1, E2, and E3 enzymes (Uba7, Ube2L6, and Herc5) have been described previously (*12*).

#### Transfections and drug treatments

HEK293T cells were plated in a 6 well plate at a density of 3×10^4^ cells/mL. The next day cells were transfected using X-tremeGENE HP DNA transfection reagent (Roche). Plasmids were transfected at the following amounts: 0.5 µg FLAG-Ubiquitin, 0.35 µg FLAG-ISG15, 0.25 µg Uba7, 0.25 µg Ube2L6, 0.35 µg HA-Herc5, 1 µg TAP-nsp123 (WT or CA), 0.2 µg TAP-PLpro^CA^, and TAP-PLpro^WT^ (0.2, 0.4, 0.6, 0.8, or 1.0 µg). Cells were treated with 6-TG at the indicated concentrations immediately following transfection (100 mM 6-TG stock in DMSO and is diluted to working concentrations in PBS).

#### Transwell assay and IL-6 ELISA

HEK293T cells (1×10^5^ cells per well) were plated in the lower chambers of 24-well Corning transwell plates (0.4 μm membrane; Fisher Scientific). At the same time, PBMCs (1×10^6^ cells per ml) were plated in a separate 10 cm dish in RPMI 1640 with 10% FBS and 1% penicillin streptomycin and allowed to rest for 16 hours. HEK293T cells were transfected with 100 ng of plasmid expressing FLAG-ISG15, with or without ISG15 conjugation components (50 ng Ube1L, 50 ng UbcH8, 100 ng Herc5) and/or SARS-CoV2 PL^pro^, using X-tremeGENE HP DNA transfection reagent (Roche) and treated with and without 0.5 µM 6-TG. 0.3 mL of PBMCs (at 1×10^6^ per mL) were added to the upper chamber of the transwell chamber, and cultured for 48 hours. Cell culture supernatants were collected from the upper transwell chamber and assayed for IL-6 production by IL-6 ELISA according to the manufacturer’s protocol. (Thermo Scientific).

#### Protein purification

ISG15 and ISG15-precursor (ISG15-pre) proteins were purified as GST fusion proteins in BL21 E. coli. Overnight starter cultures were grown at 37°C for 16 hours. Cultures were diluted 1:10 and cultured with shaking for 2 hours at 37°C. Expression of proteins was induced with 100 µM Isopropyl β-D-1-thiogalactopyranoside (IPTG) for 3 hours at 30°C. Cells were collected by centrifugation, resuspended in 10 mL PBS with 1% Triton X-100, and sonicated for 30 seconds. Lysates were spun at 15,000xg for 10 minutes, and supernatants were incubated with 100 µL Glutathione Sepharose (GE Healthcare) for 2 hours with rotation at 4°C. Beads were collected and washed 3X with PBS plus 1% Triton X-100 and 3X with 50mM Tris pH 8.0, 150mM NaCl, 0.01% Triton X-100. GST-ISG15-pre was eluted in 100µL of 10 mM reduced GSH for 1 hr at room temperature.

Sars-CoV-2 PLpro was purified as a GST fusion protein in BL12 E. coli. Overnight starter cultures were grown at 37°C for 16 hours. Cultures were diluted 1:10 and cultured with shaking for 2 hours at 37°C. Expression of proteins was induced with 100 µM Isopropyl β-D-1-thiogalactopyranoside (IPTG) for 24 hours at 16°C. Cells were collected by centrifugation, resuspended in 10 mL TBS with 0.00025% Tween 20, 0.01 mM DTT and 5% glycerol (Binding Buffer), and sonicated for 1 minute. Lysates were spun at 15,000xg for 10 minutes and supernatants were incubated with 100 µL per liter of culture Glutathione Sepharose (GE Healthcare) for 4 hours with rotation at 4°C. Beads were collected and washed three times with Binding Buffer supplemented with 0.1mM EDTA and subjected to site-specific cleavage with PreScission Protease (GE Healthcare) to remove the GST tag. Beads were removed and the protein concentration in the supernatant was quantified by SDS-PAGE using a Licor Odyssey Imager.

#### In vitro cleavage of GST-ISG15-pre

0.1 µM PLpro was incubated with 0.1 µM GST-ISG15-pre for the indicated amount of time with the indicated amount of 6-TG. Reactions were initiated by the addition of GST-ISG15-pre and stopped by addition of 10µL of 250mM tris pH 6.8, 40% glycerol 8% SDS and 400mM DTT. Samples were run on 10% Tris-Glycine gels and stained with Coomassie blue stain. Gels were imaged using a Licor Odyssey Imager.

#### Immunoblotting and immunoprecipitations

Samples assessing ISGylation or ubiquitination were lysed in 1% NP40 lysis buffer (100 mM Tris pH 7.5, 100 mM NaCl,1% NP40) supplemented with 10 mM NEM, 170 µg/mL PMSF, 2 µg/mL leupeptin, 2 µg/mL aprotinin and 10 mM DTT for 10 minutes on ice followed by a 10 minute spin at 20000 x *g*. Samples were run on BOLT 4-12% Bis-Tris gels (Thermo) and blotted with the M2 FLAG antibody (Sigma). Samples assessing the effect of 6-TG on TAP-nsp123 cleavage were lysed in RIPA buffer (50 mM Tris pH 8, 150 mM NaCl, 0.5% NP40, 0.1% SDS, 0.5% Sodium deoxycholate w/v) supplemented with 10 mM NEM, 170 µg/mL PMSF, 2 µg/mL leupeptin, 2 µg/mL aprotinin, 10 mM DTT, and 2 units/mL Universal Nuclease (Pierce) for 10 minutes on ice followed by a 10 minute spin at 20000 x *g*. Samples were run on 3-8% Tris-Acetate gels (Thermo) and blotted for TAP using the anti-protein A antibody (Sigma). Actin blots were performed with an anti actin antibody clone: AC-15 (Invitrogen).

For detection of extracellular ISG15, cell culture supernatants were collected and cleared of any cell debris by centrifugation at 1000 x *g*. The supernatants were further cleared of IgG from the serum by two one-hour bindings with Protein A sepharose (Invitrogen) at room temperature. 20 µL of Sera-Mag SpeedBead Protein A/G magnetic beads (GE Life Sciences), 10 µL of ISG15 antibody (Invitrogen 7H29L24) and 20 mL of 0.1% NP40 in PBS pH 7.0 were added to the cell culture supernatants and rocked for 2 hours at room temperature. Beads were isolated with a neodymium magnet and washed 3X with 0.1% NP40 in PBS pH 7.0. ISG15 was eluted from the beads by boiling in 80 µl of 1X loading buffer and 40 µl was run on a NuPAGE 4-12% Bis-Tris gel and transferred to nitrocellulose for western blotting. ISG15 was detected with the M2 Flag antibody (Sigma).

#### Virus replication assays

1.5-2 x 10^5^ Vero-E6 and 5 x 10^5^ Calu3 cells were seeded in a 24 well format. The cells were infected with SARS-CoV-2 at a MOI of 0.001 diluted in culture media containing 2% FBS. Virus inoculum was removed after one hour and cells were washed with 1mL PBS twice. Infected cells were cultured in media containing 6-TG (25 µM, 5 µM, 1 µM, 200nM, 40nM, 8nM, NoTx). 100 µl of culture supernatants were harvested into (ORF) 300 μL TRK lysis buffer plus BME at 24 and 48 hours. RNA was isolated from homogenates of TRK lysates using an E.Z.N.A. Total RNA kit I (Omega). Isolated RNA was resuspended in UltraPure H_2_O (Ambion). RT-qPCR was performed on a ViiA 7 or an ABI 7500 qPCR machine (Thermo Scientific) using the TaqMan RNA-to-CT 1-Step Kit (Applied Biosystems) with a 20 μL total reaction volume per well including 8.5 μL of RNA sample and Taqman primers and SARS-CoV2 N1 probe of 2019-nCoV RUO Kit from Integrated DNA Technologies. The following PCR cycling protocol was used: 15 minutes at 48°, 10 min at 95°, and 50 cycles of 15 sec at 95° and 1 min at 60°C. RNA copy number was determined for each sample using the matching RNA standard (gift of Adam Bailey, Michael S. Diamond lab, Washington University School of Medicine).

#### 6-TG toxicity assays

The real-time cytotoxicity events were analyzed by using a 2-color IncuCyte ZOOM in-incubator imaging system (Essen Bioscience). Dead cells were detected by uptake of the cell-impermeable dye SYTOX Green (Invitrogen). The total cell numbers were estimated by calculating cell numbers staining with cell-permeable dye SYTO24.

#### Quantification and Statistical analysis

Graphpad Prism 8 software was used to plot ELISA, virus replication and densitometry data. Bars represent three or more biological replicates. Error bars represent the standard error of the mean (SEM). Ordinary one-way ANOVA was performed on each ELISA data set comparing all conditions to treatment with 6-TG, alone. EC_50_ and CC_50_ of 6-TG were calculated using “log(inhibitor) vs. normalized response -- Variable slope”. Cell viability, dose response curve, and EC_50_ with 95% confidence interval was plotted/indicated in the graph. Asterisks indicate p-values: *=0.02, **=0.002, ***=0.004, ****<0.0001, and non significant changes are indicated by ns.

**Fig. S1.**
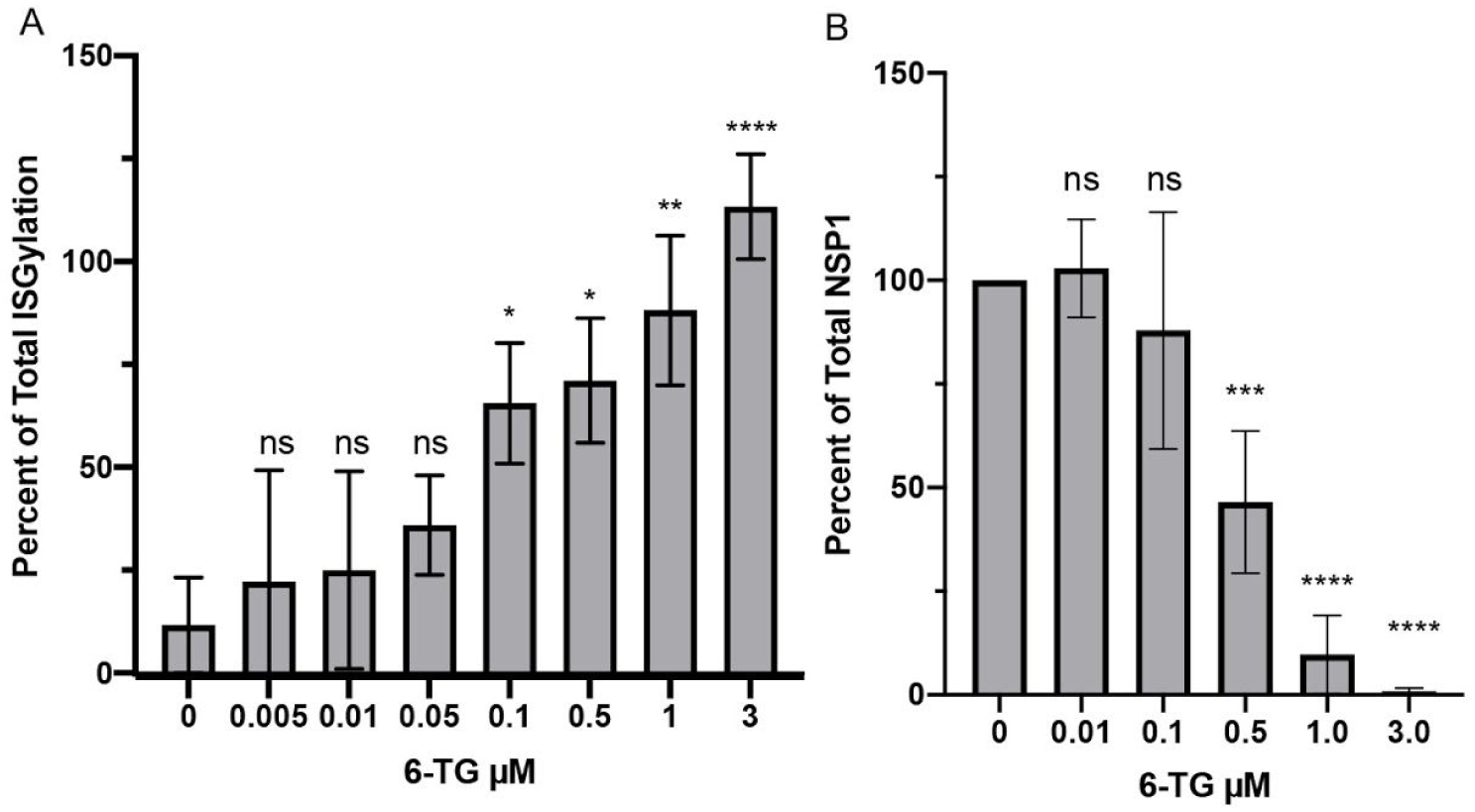
Quantification of ISGylation and nsp123 cleavage by PLpro. **A.** Densitometry analysis of ISGylation from Fig. 1A. Percent of total ISGylation was determined by comparison to total ISGylation in the presence of PLpro^CA^. **B.** Densitometry analysis of TAP-nsp1 from Figure 1B. Percent TAP-nsp1 was determined by comparison to the 42 kDa band in the untreated samples to those present in samples treated with 6-TG. Significance was assessed by ordinary one-way ANOVA comparison to treatment with no 6-TG. Asterisks indicate p-values: *=0.02, **=0.002, ***=0.004, ****<0.0001, and non-significant changes are indicated by ns.

